## Supplemental Figures for "TPGS1 Regulates Central Spindle Microtubule Glutamylation and Remodeling During Telophase and Abscission"

### Supplementary Figure 1.

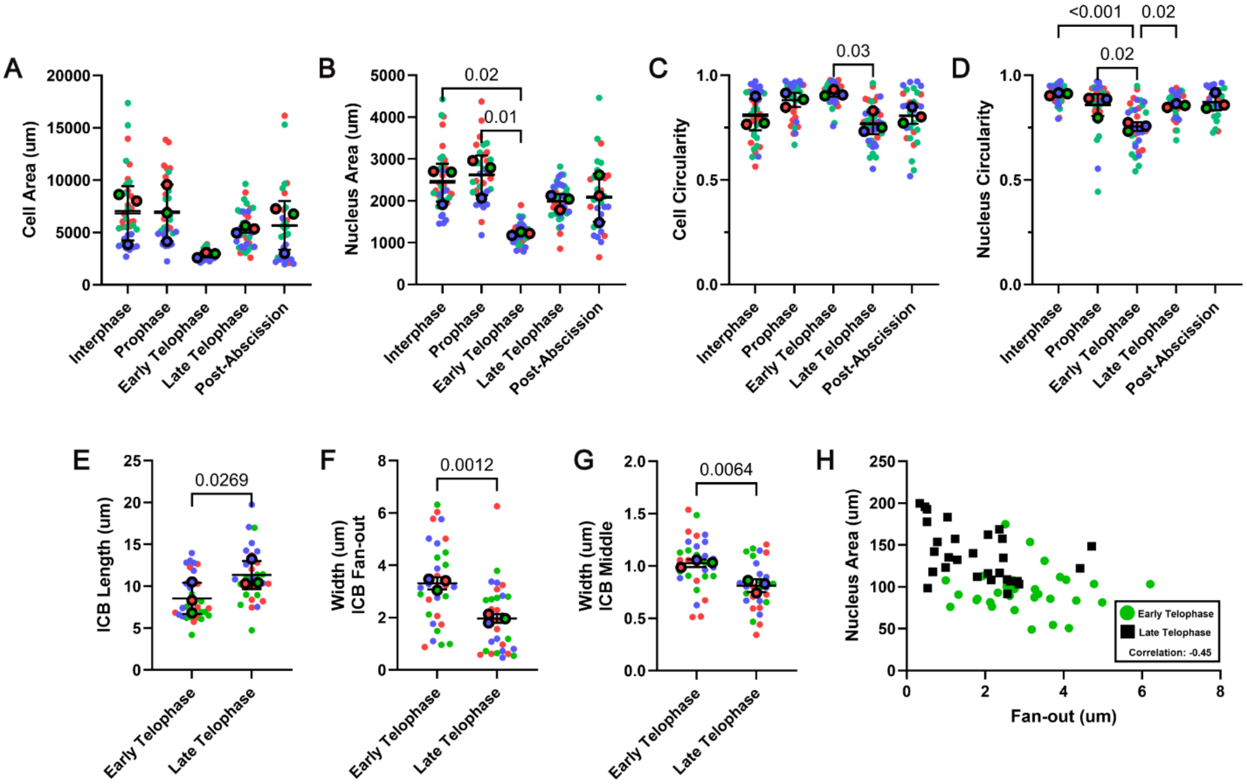

**Supplementary Figure 2.**

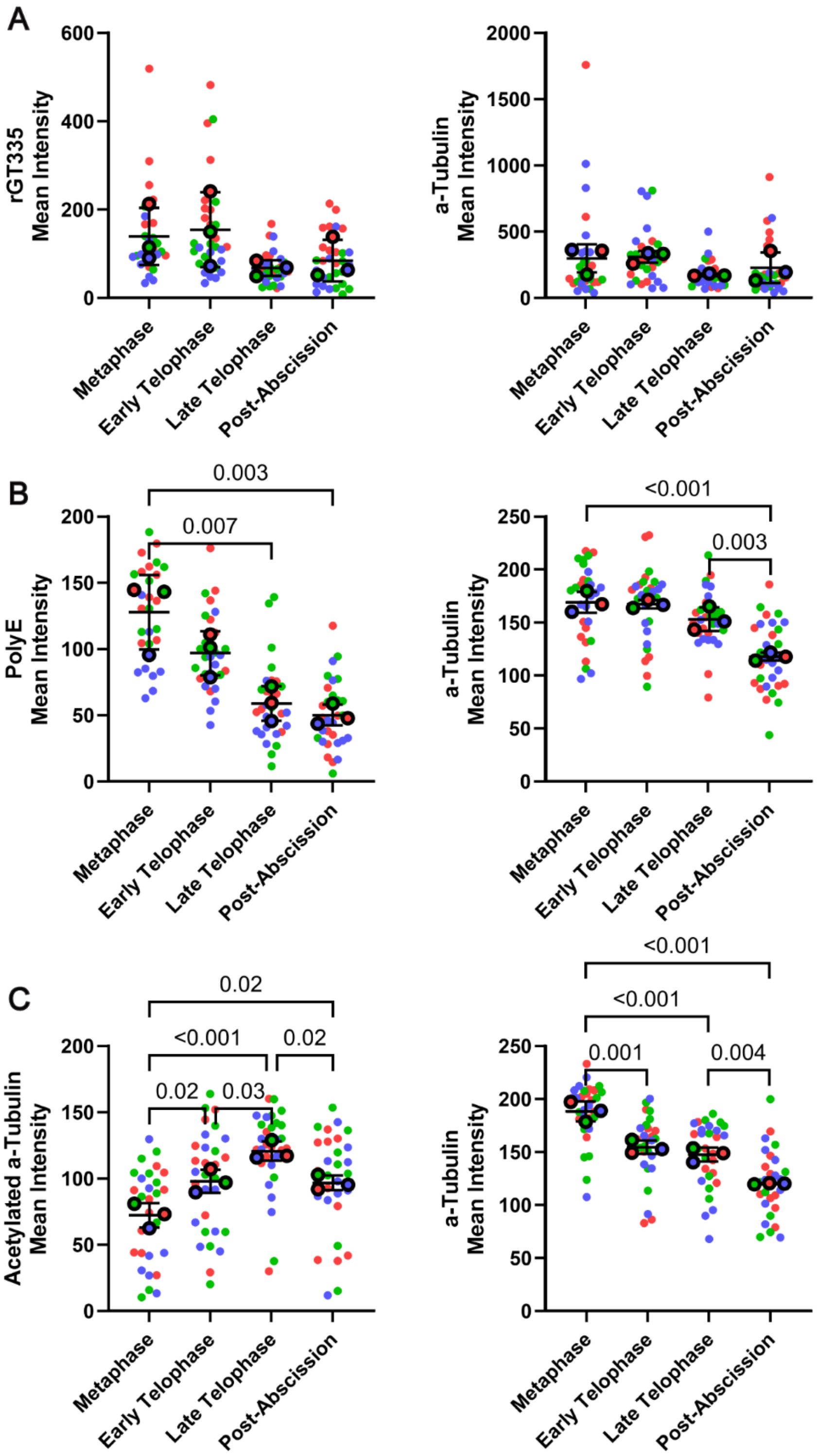

**Supplementary Figure 3.**

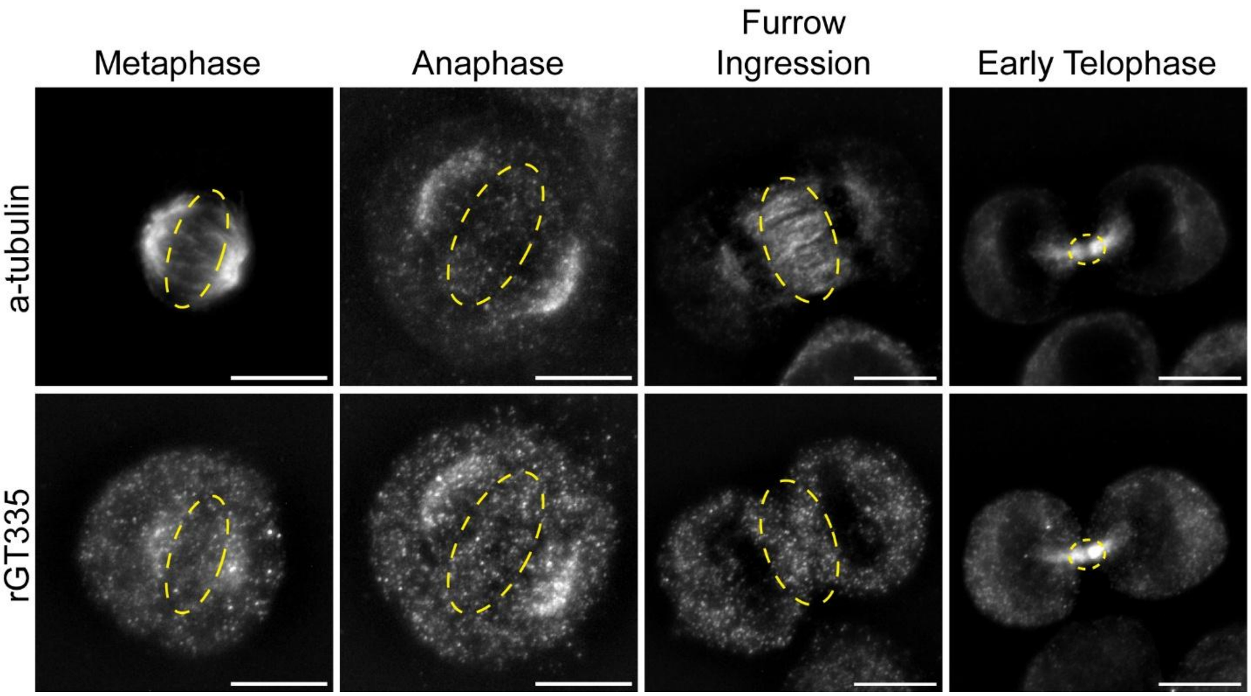

**Supplementary Figure 4.**

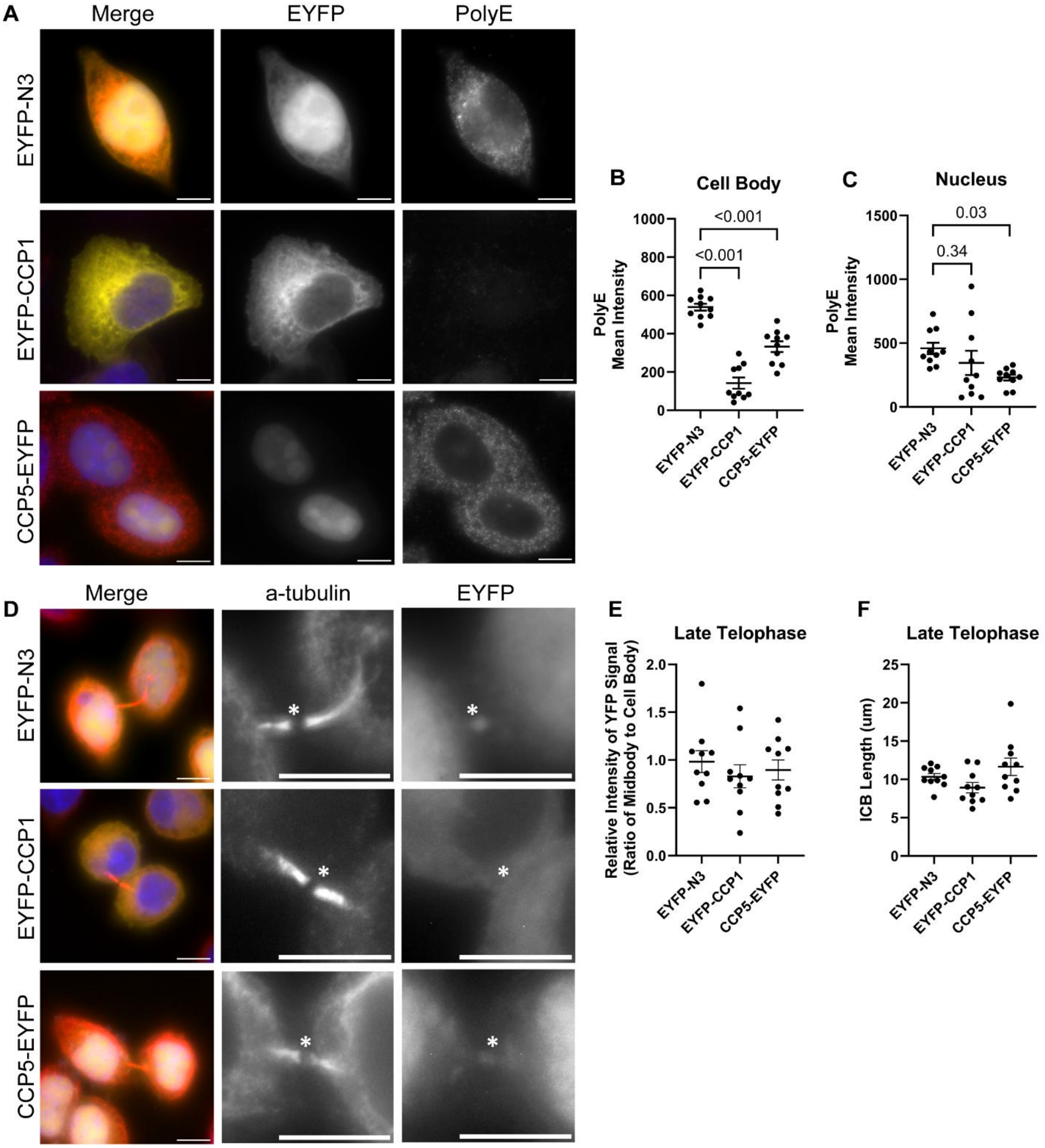

**Supplementary Figure 5.**

**A**

```
                575                                                                              656
TPGS1 Gen...  CCGCCGGCCGGTTTCACGGACAGCGGCCGCCAGTCGGTATCCCGGGCGGCGGGGGCGGCCGAGAGCGAGGAGGACTTCCTGC
10.1_TPGS...  CCGCCGGCCGGTTTCACGGACAGCGGCCGCCAGTCGGTATCCCGGGCGGCGGGGGCGGCCGAGAGCGAGGAGGACTTCCTGC
TPGS1-KO-...  CCGCCGGCCGGTTTCACGGACAGCGGCCGCCAGTCGGTATCCCGGGCGGCGGGGGCGGCCGAGAGCGAGGAGGACTTCCTGC
TPGS1-KO-...  CCGCCGGCCGGTTTCACGGACAGCGGCCGCCAGTCGGGATCCCGGGCGGCGGGGGCGGCCGAGAGCGAGGAGGACTTCCTGC
TPGS1-KO-...  CCGCCGGCCGGTTTCACGGACAGCGGCCGCCAGTCGGTATCCCGGGCGGCGGGGGCGGCCGAGAGCGAGGAGGACTTCCTGC
TPGS1-KO-...  CCGCCGGCCGGTTTCACGGACAGCGGCCGCCAGTCGGTATCCCGGGCGGCGGGGGCGGCCGAGAGCGAGGAGGACTTCCTGC
TPGS1-KO-...  CCGCCGGCCGGTTTCACGGACAGCGGCCGCCAGTCGGTATCCCGGGCGGCGGGGGCGGCCGAGAGCGAGGAGGACTTCCTGC
TPGS1-KO-...  CCGCCGGCCGGTTTCACGGACAGCGGCCGCCAGTCGGTATCCCGGGCGGCGGGGGCGGCCGAGAGCGAGGAGGACTTCCTGC

..............................................................................................

                657                                                                              738
TPGS1 Gen...  GGCAGGTCGGCGTGACGGAAATGCTACGTGCGGCCCTGCTGAAGGTGCTGGAGGCGCGGCCCGAGGAGCCGATCGCCTTCCT
10.1_TPGS...  GGCAGGTCGGCGTGACGGAAATGCTACGTGCGGCCCTGCTGAAGGTGCTGGAGGCGCGGCCCGAGGAGCCGATCGCCTTCCT
TPGS1-KO-...  GGCAGGTCGGCGTGACGGAAATGCTACGTGCGGCCCTGCTGAAGGTGCTGGAGGCGCGGCCCGAGGAGCCGATCGCCTTCCT
TPGS1-KO-...  GGAAGGTCGGTGTGACGGAAATGCTACGTGCGGCCCTGCTGAAGGTGCTGGAGGCGCGGCCCGAGGAGCCGATCGCCTTCCT
TPGS1-KO-...  GGCAGGTCGGCGTGACGGAAATGCTACGTGCGGCCCTGCTGAAGGTGCTGGAGGCGCGGCCCGAGGAGCCGATCGCCTTCCT
TPGS1-KO-...  GGCAGGTCGGCGTGACGGAAATGCTACGTGCGGCCCTGCTGAAGGTGCTGGAGGCGCGGCCCGAGGAGCCGATCGCCTTCCT
TPGS1-KO-...  GGCAGGTCGGCGTGACGGAAATGCTACGTGCGGCCCTGCTGAAGGTGCTGGAGGCGCGGCCCGAGGAGCCGATCGCCTTCCT
TPGS1-KO-...  GGCAGGTCGGCGTGACGGAAATGCTACGTGCGGCCCTGCTGAAGGTGCTGGAGGCGCGGCCCGAGGAGCCGATCGCCTTCCT

..............................................................................................

                739                                                                              820
TPGS1 Gen...  GGCTCACTACTTCGAGAACATGGGCCTGCGCTCGCCTGTAAACGGCGGCGCCGGGGAGCCCCCGGGCCAGCTCCTGCTGCAG
10.1_TPGS...  GGCTCACTACTTCGAGAAGACGG----------GCCTGTAAACGGCGGCGCCGGGGAGCCCCCGGGCCAGCTCCTGCTGCAG
TPGS1-KO-...  GGCTCACTACTTCGAGAAGACGG----------GCCTGTAAACGGCGGCGCCGGGGAGCCCCCGGGCCAGCTCCTGCTGCAG
TPGS1-KO-...  GACTCACTACTTCGAGAAAACGG----------GCCTGTAAACGGCGGCGCCGGGGAGCCCCCGGGCCAGCTCCTGCTGCAG
TPGS1-KO-...  GGCTCACTACTTCGAGAAGACGG----------GCCTGTAAACGGCGGCGCCGGGGAGCCCCCGGACCAGCTCCTGCTGCAG
TPGS1-KO-...  GGCTCACTACTTCGAGAAGACGG----------GCCTGTAAACGGCGGCGCCGGGGAGCCCCCGGGCCAGCTCCTGCTGCAG
TPGS1-KO-...  GGCTCACTACTTCGAAAAGACGG----------GCCTGTAAACGGCGGCGCCGGGGAGCCCCCGGGCCAGCTCCTGCTGCAG
TPGS1-KO-...  GGCTCACTACTTCGAGAAGACGG----------GCCTGTAAACGGCGGCGCCGGGGAGCCCCCGGGCCAGCTCCTGCTGCAG
```

**B**

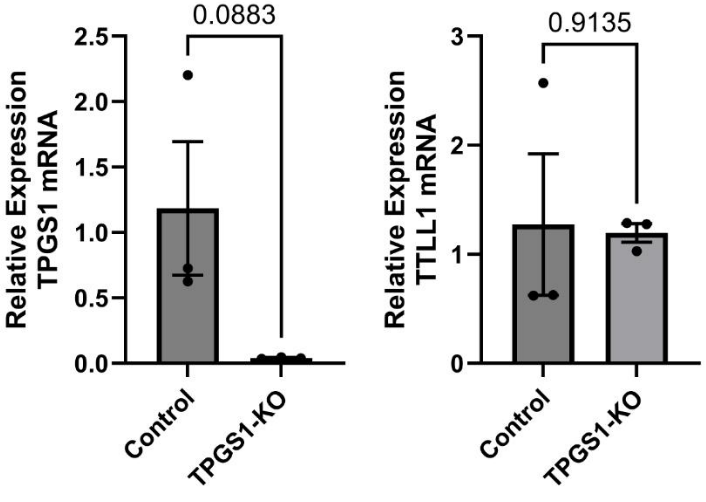

**Supplementary Figure 6.**

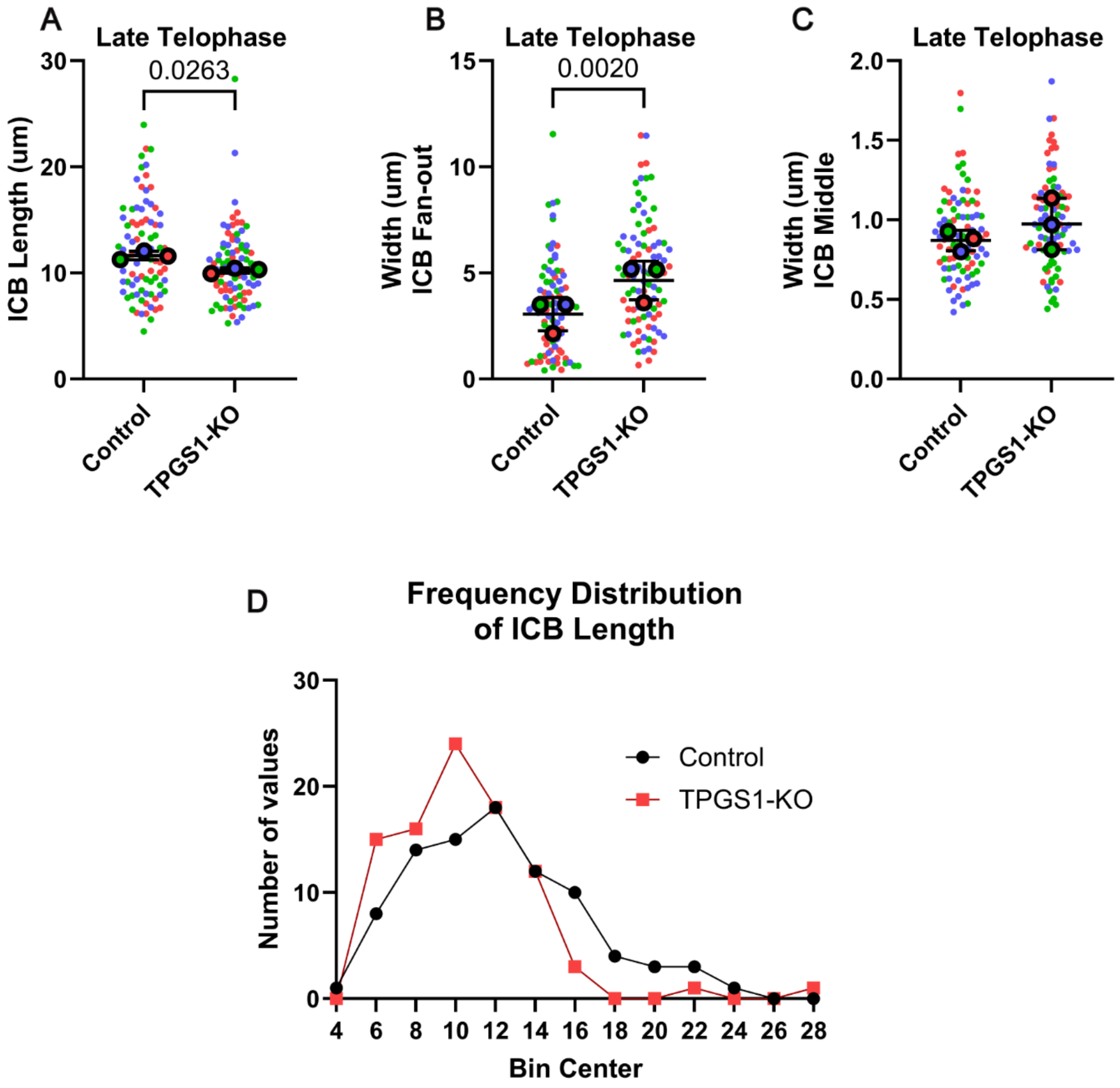

**Supplementary Figure 7.**

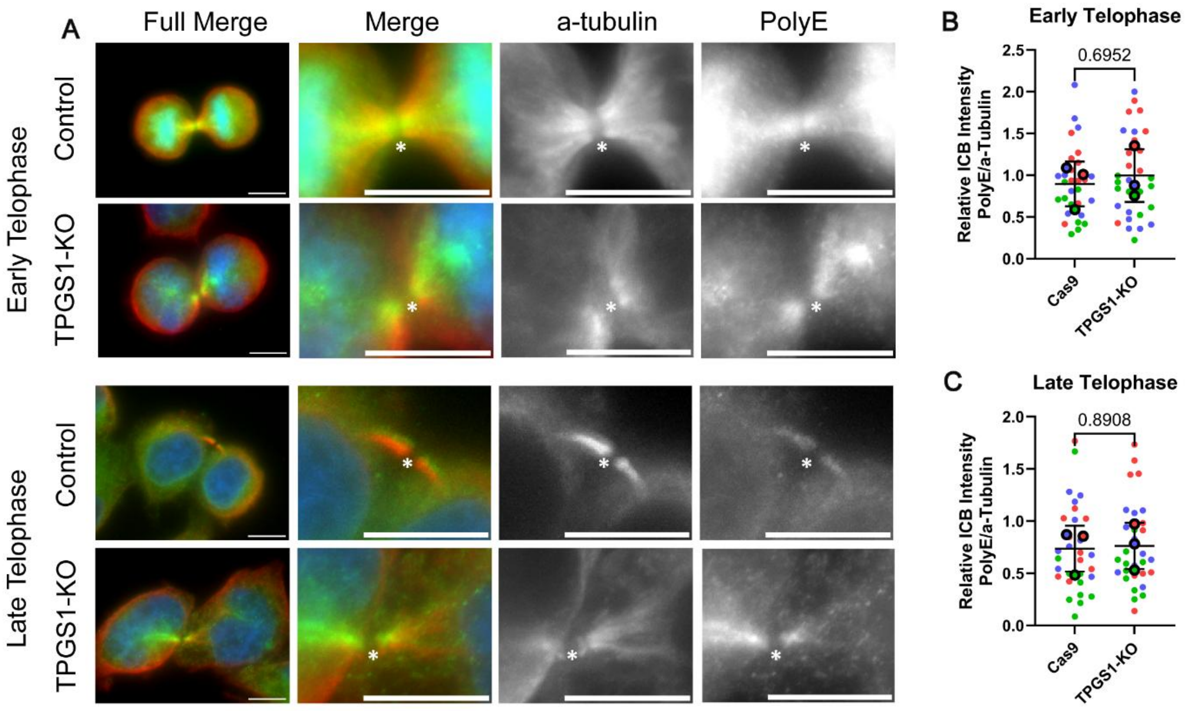

**Supplementary Figure 8.**

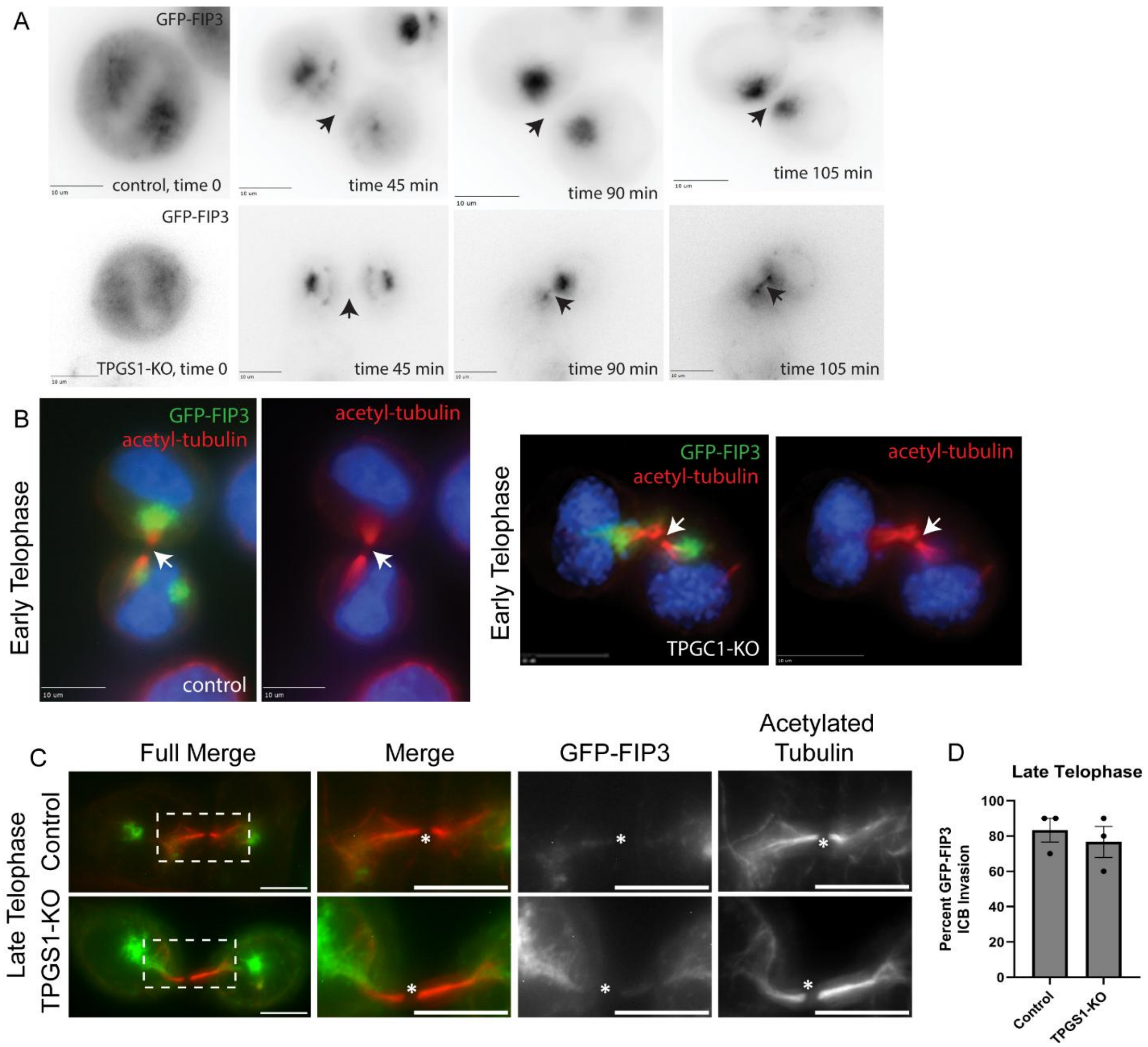

**Supplementary Figure 9.**

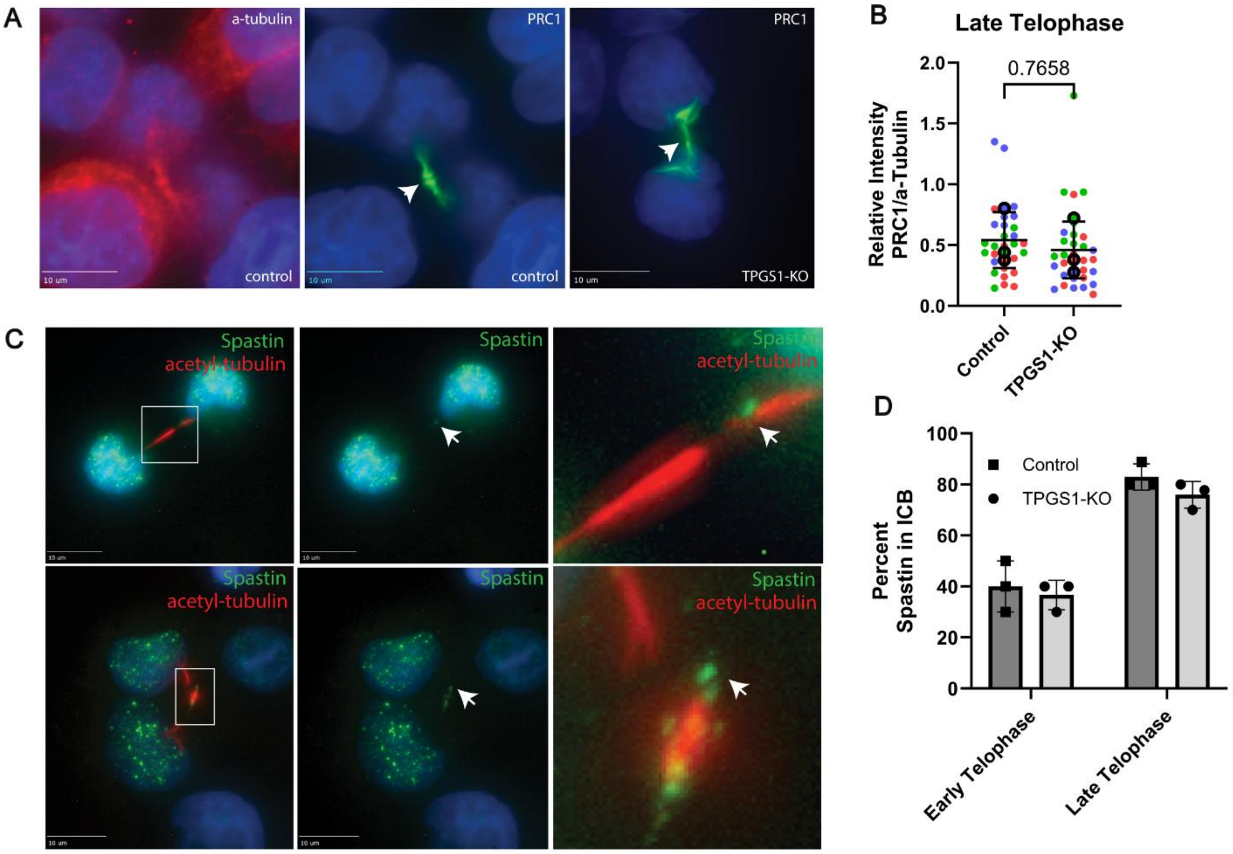

#### SUPPLEMENTAL FIGURE LEGENDS

##### Supplemental Figure 1.

(A-H) Quantification of images of fixed wild-type HeLa cells, co-stained with and anti-$\alpha$-tubulin and anti-acetylated $\alpha$-tubulin antibodies. Each experiment included biological replicates (=3), with 10 cells measured per replicate, color coded. One-way ANOVA was done on (A-D), t-test on (E-G), and linear regression analysis on (H).

(A-D) Cell and nucleus area and circularity were measured by selecting $\alpha$-tubulin signal (cells; A,C) and Hoechst signal (nuclei; B and D) to measure total area and circularity.

(E-G) Telophase cell ICB dimensions used for ratios in figure 1 were measured between each pair of cells, each treated as one technical replicate per one pair of cells.

(H) Nucleus area plotted relative to paired fan-out value, indicating a negative correlation between the two. Nucleus area was a primary indicator of telophase stage used in this paper. Early telophase cells are shown in green and late telophase cells in black.

##### Supplemental Figure 2.

(A-C) Mean fluorescence intensity values of microtubule structures used for ratios shown in figure 2, given for each antibody pair: anti-rGT335 (A), anti-PolyE (B), and anti-acetylated $\alpha$-tubulin (C). Each antibody was co-stained with anti-$\alpha$-tubulin antibodies. Statistics were calculated with one-way ANOVA on means from color-coded biological replicates (=3), each including 10 cells.

##### Supplemental Figure 3.

Representation from figure 3 of where masks were drawn for calculation of Pearson's and Mander's correlations. Masks incorporate the area containing the anti-parallel overlap of central spindle microtubules. Image scale bars are 10$\mu$m.

**Supplemental Figure 4.**

(A-C) Representative images and quantification of cells transfected with EYFP-N3, EYFP-CCP1, and CCP5-EYFP plasmids (yellow), fixed and co-stained with anti-PolyE (red) antibodies. Quantification of PolyE signal was done on 10 cells in one biological replicate, measured for the cell body (B) (with the nucleus subtracted) and the nuclus (C). Statistics were done with one-way ANOVA, but only represent that of technical replicates. Image scale bars are 10μm.

(D) Images of telophase cells transfected with EYFP-N3, EYFP-CCP1, and CCP5-EYFP plasmids (yellow), fixed and co-stained with anti-α-tubulin (red) antibodies. Image scale bars are 10μm. Asterisk marks the MB.

(E) Ratio of EYFP fluorescence mean intensity of the MB divided by that of the whole cell per image, of one biological replicate containing 10 cells in each condition.

(F) ICB length measured between telophase cells in each condition of one biological replicate containing 10 cells in each condition.

**Supplemental Figure 5.**

(A) Sequencing results aligned to TPGS1 sequence for clones of TPGS1-KO cells. Cells exhibit a 10bp deletion in all alleles resulting in frame shift and premature stop codon.

(B) RT-qPCR results on biological replicates (=3) of control and TPGS1-KO cells for TPGS1 and TTLL1 mRNA, normalized to GAPDH. Error bars represent SEM.

**Supplemental Figure 6.**

(A-D) Quantification of images of fixed wild-type HeLa cells, co-stained with and anti-α-tubulin and anti-acetylated α-tubulin antibodies. Each experiment included biological replicates (=3), with 30 cells measured per replicate, color coded. All statistics are done on means. T-test was calculated for (A-C), Late telophase cell ICB dimensions of control and TPGS1-KO cells used for ratios in figure 7 were measured between each pair of cells, each treated as one technical replicate per one pair of cells.

**Supplemental Figure 7.**

(A-C) Imaging and quantification of early telophase (A-B) and late telophase (A,C) control and TPGS1-KO HeLa cells. Cells were fixed and co-stained with anti-PolyE (green) and anti-$\alpha$-tubulin (red) antibodies. All image scale bars are 10$\mu$m. Asterisk marks the MB. Biological replicate (=3) means are color coded, with 10 cells per replicate. Statistics are calculated with t-test on mean values for each stage. Error bars represent the SEM surrounding mean values.

**Supplemental Figure 8.**

(A) Time-lapse imaging of control and TPGS1-KO HeLa cells transfected with GFP-FIP3. Scale bars are 10$\mu$m, arrows point to the MB.

(B-C) Control and TPGS1-KO cells were transfected with GFP-FIP3 (green), fixed, and stained with anti-acetylated-$\alpha$-tubulin (red) antibodies. Arrows in (B) and asterisks in (C) mark the MB. Scale bars are 10$\mu$m. Boxes mark the zoomed in area.

(D) Quantification of images from (C) of late telophase cells, measuring the percent of cells in each biological replicate (=3) that showed GFP-FIP3 signal in the ICB. T-test was done on the percentages of replicates.

**Supplemental Figure 9.**

(A) Control and TPGS1-KO cells were fixed and stained with anti-$\alpha$-tubulin (red) and anti-PRC1 (green) antibodies. Arrows mark the MB. Scale bars are 10$\mu$m.

(B) Relative intensity of PRC1 signal normalized to $\alpha$-tubulin in late telophase of cells shown in (A). Biological replicate means (=3) are color coded and statistics were done with t-test of the means, with each mean representing 10 cells.

(C) Control cells were fixed and stained with anti-acetylated $\alpha$-tubulin (red) and anti-spastin (green) antibodies. Arrows mark spastin localization at microtubule severing sites. Scale bars are 10$\mu$m. Boxes mark the zoomed in area.

(D) Quantification of images from (C) and of TPGS1-KO cells in early and late telophase, measuring the percent of cells in each biological replicate (=3) that showed spastin localized near the MB in intact ICBs in images of cells. T-test was done on the percentages of replicates between each stage of telophase.
